# Expanding perspectives on cognition in humans, animals, and machines

**DOI:** 10.1101/037192

**Authors:** Alex Gomez-Marin, Zachary F Mainen

**Affiliations:** Champalimaud Neuroscience Programme, Champalimaud Centre for the Unknown, Lisbon, Portugal.

## Abstract

Over the past decade neuroscience has been attacking the problem of cognition with increasing vigor. Yet, what exactly *is* cognition, beyond a general signifier of anything seemingly complex the brain does? Here, we briefly review attempts to define, describe, explain, build, enhance and experience cognition. We highlight perspectives including psychology, molecular biology, computation, dynamical systems, machine learning, behavior and phenomenology. This survey of the landscape reveals not a clear target for explanation but a pluralistic and evolving scene with diverse opportunities for grounding future research. We argue that rather than getting to the bottom of it, over the next century, by deconstructing and redefining cognition, neuroscience will and should expand rather than merely reduce our concept of the mind.

## 1. Introduction

Neuroscience aims to explain how the workings of the mind and behavior arise from the biology of interacting cells and molecules. This is arguably the biggest scientific challenge of our era. One of the main reasons that this problem is so hard is obvious: it is the vast complexity of the brain. But a somewhat less obvious part of the problem is agreeing on the question: what exactly are we trying to explain? The term “cognition” is used in neuroscience as shorthand to refer to the most complex functions the brain enables^1^. Cognition has traditionally been defined in terms of a suite of “cognitive functions”—attention, working memory, and so on. But are these really the key building blocks for complex behavior? If not, the definition of cognition should be realigned on a better-grounded empirical or theoretical approach. But where is that to be found? These questions are critical because without a firm grasp of what exactly is cognition, apparent progress in understanding its neural basis may evaporate as it is realized that the wrong object was under scrutiny.

In this review we survey contemporary approaches to brain complexity that might serve as a basis for redefining cognition. We begin by very briefly reviewing over 40 years of progress in studying the single neuron correlates of cognition. We emphasize the recent theoretical proposal that the complexity of these cognitive tasks is extremely low, implying that we might indeed be missing something in the definition. From this we launch into a broader discussion of possible theoretical bases for defining cognition. We then turn to consider the more empirical bases found in machine learning, behavioral studies, and neurophenomenology.

## 2. Neural basis of cognition in monkeys and rodents

The idea of cognition goes back at least to Aristotle (*gignoskein*, to know, the process of acquiring knowledge) but didn’t really get going in a scientific form until the 19th century. The field of psychology has sought for many years to understand the basis of intelligence—cognition being the name for the higher capacities of the mind. By the time of Ebbinghaus and James in the late 19th century, many forms of cognition were beginning to be subjected to scientific study by psychologists.

Starting in the 1960’s neuroscientists began to translate work in cognitive psychology from humans into monkeys. The goal of this work was to reveal direct physical correlates of cognitive functions such as working memory^2^. To do this, they systematically recorded the electrical activity of single neurons in behaving animals. This required them to teach monkeys (macaques) to perform cognitive tasks. This line of work demonstrated that monkeys not only exhibit behaviors considered to constitute the most basic building blocks of cognition— learning & memory, decision-making—but can perform more complex tasks including selective attention^3^, flexible categorization^4^, integration of evidence^5^, probabilistic reasoning^6^, metacognitive judgements^7^, numerical reasoning^4^, context-dependent decisions^8^ and more.

Starting in the 1980’s, researchers began to show that even rodents can also perform a wide variety of standard cognitive tasks including selective attention^9^, flexible sensorimotor mapping^10^ and working memory. In the last decade or so, this line of work in freely-behaving animals was refined to even more exacting standards, allowing quantitative measurements of single neuron activity in relationship to functions including confidence judgments^11^, psychophysical thresholds^12^, integration of evidence^13^, self-initiated actions^14^, cue combination^15^, and flexible sensory-motor associations^16^. A burgeoning line of work has taken this approach to even more constrained, head-fixed, mice performing tasks such as memory guided decisions^17^, sometimes in a virtual reality^18^.

## 3. Cognition in just a few dimensions

This work has set the stage for huge progress in “opening the black box”, as it allows rodents—which are smaller, cheaper, far easier to experiment with, less ethically demanding—to stand in for monkeys or even humans. Particularly because this allows the use of molecular and genetic manipulations, which have become more and more powerful^19^,^20^, many researchers using macaques have begun to switch to rats or mice or smaller, faster-breeding marmosets, in which genetic manipulations may be performed with greater ease.

Yet the desire to account for behavior on the basis of single spikes in single neurons raises a rather perplexing issue. It seems preposterous to hope to understand the concerted actions of a brain of 100 million neurons by studying a handful of neurons at a time. Yet, somehow this does not seem to be the case. Single neurons or populations <100 have typically revealed meaningful correlates of target cognitive phenomena. When analyzed using dimensionality reduction techniques, even the most complex tasks mentioned have yielded a dimensionality that is far lower than even the relatively small number of neurons being analyzed^8^,^21^.

What is going on? Ganguli and colleagues have proposed a theory of “neural task complexity”^22^. They argue that as long as task parameters are limited and neuronal activity patterns vary smoothly across task parameters, that even for what we think of as complex cognitive tasks, the neural task complexity is very small. This work suggests that rather than recording more neurons, in order to make progress we need richer, more complex tasks^22^. This relates to the important issue of how much of what we measure in behaving animals is due to behavior itself or simply reflecting the environment in which the animal is placed in the laboratory. It also raises the more radical possibility that neuroscience may already be outgrowing the traditionally defined set of cognitive functions.

## 4. A basis in psychology or in the brain itself?

Nearly 80 years ago, Tolman proclaimed that “everything important in psychology (⃛) can be investigated in essence through the continued experimental and theoretical analysis of the determiners of rat behavior at a choice-point in a maze”^23^. Our current cognitive taxonomy appears to include a much richer set of behaviors, but much of our basic cognitive toolkit was already old hat to Tolman, having been developed decades ago by psychologists such as William James^24^. Granting James’ genius, how much confidence can be taken that the 19^th^ century agenda of cognitive psychology covers the breadth and depth of mental function?

The idea of dissecting cognition into isolatable components, combined with the possibility of studying the neural basis of such a taxonomy of sub-functions (attention, working memory, meta-cognition, etc.) appears to be well-suited to the paradigm set by molecular biology and genetics for the study of biological mechanism. The prominence of neural reductionism, “the tendency to seek all causal relations of behavior in brain processes”^25^, has caused some to wonder whether psychological explanation is in danger of extinction^26^. This is a bias that might be exacerbated in the current climate of increased competition for publication and funding. What does it mean to explain cognition^27^? If explaining entails explaining it away (if making it clearer results in dissolving it into low level mechanisms) then the drive toward reductionistic accounts may ironically turn into a roadblock in our understanding. Instead, here we attempt to redefine cognition from its multiple approaches. Truly novel work does go on at the psychological level, even if it has a harder time finding an audience without a neural substrate. For instance, while Bayesian-inspired psychological research has made inroads into mechanistic cognitive neuroscience, cognitive psychologists continue to work beyond what is cutting edge in neuroscience on problems such as rapid structural learning and concept learning^28^.

It has been argued that psychology suffers from human-invented categories that do not necessarily conform to nature—“we take a man-created word or concept (⃛) and search for brain mechanisms that may be responsible for the generation of this conceived behavior”^29^. An alternative is an inside-out or bottom-up approach building from brain mechanisms toward cognitive function. While this strategy offers freedom from potentially awkward hypotheses about the phenomena to be explained, for a system as complex as a brain, a pure bottom up approach appears to be a non-starter. The power of discovery based on sophisticated techniques relying on big data can fail to produce significant results but, in a sense, it can never be wrong. Therefore falsifiable theories and concrete testable hypothesis — not just impressive tour-de-force— are needed. Even with an organism for which the complete wiring diagram is available, *C. Elegans,* behavioral function cannot be deduced without additional hypotheses. This problem is also illustrated by the heroic but ultimately humbling attempt to model a single cortical column from the bottom-up^30^.

## 5. Theories and frameworks for cognition

If one is to approach and define cognition at least partly from the top-down, what are the available theoretical frameworks from which this might be done? Here we find several alternatives, not necessarily mutually exclusive. Such theoretical approaches are rooted in mathematics and deep concepts—probabilistic inference, learning, entropy, complexity—that are being mined for their potential as an ultimate level of explanation for brain function.

### Dynamical systems

A long-standing theoretical approach to cognition is based on nonlinear dynamical systems theory, which has been proposed to be a noncomputational way to conceive of cognition^31^. Two claimed advantages of conceiving of a cognitive agent as a dynamical system, rather than a digital computer, are the possibility to interact with the world without “representing” it and the possibility to avoid discretizing time^32^. Dynamical systems approaches were the basis for understanding the basis of neuronal function at the cellular level^33^. They are also the basis for the more “biophysical” network models, which have demonstrated that the dynamics of very simple neuronal circuits can be turned to perform elementary cognitive functions such as working memory and decision-making^34^,^35^. Moreover, with appropriate (albeit sometimes implausibly nonlocal) learning rules these networks can be endowed with surprisingly rich dynamics, e.g. generating arbitrary dynamical mappings^36^,^37^, providing mechanistic hypothesis for the basis of context-dependent decisionmaking. Dynamical network models based on engineering rather than learning solutions have begun to be assembled to perform complete tasks^38^ but cannot say play a video game. Approaches based on optimization (learning) of relatively simple objective functions are remarkably successful in capturing the dynamics of complex actions (e.g. standing up) using nonlinear dynamical models of the motor plant and physical environment^39^. Work from this perspective argues that cognition and action are deeply intertwined, an “embodied” perspective reinforced by the loopiness of brain anatomy. Motor-control inspired dynamic systems approaches may provide fertile ground for future neuroscience of cognition.

### Reinforcement learning

Over the last two decades, reinforcement learning (RL) theory has been one of the most impactful theoretical frameworks in neuroscience^40^. With roots in artificial intelligence, to which we return below, the success of RL in neuroscience has been forged by theorists who have taken a very much top-down or normative approach to ask what *should* the brain do. Importantly, RL theorists have also cared deeply about the underlying brain circuitry, particularly at the systems level (area by area), and the behavior, particularly that carried out by the “classical” animal learning theorists (essentially the line of Skinner). Although the name emphasizes one flavor of learning, RL naturally incorporates decision-making, and strong relationships with microeconomic theory exist^41^. RL deals with optimization based on an externally-supplied scalar objective function. Yet it should not be forgotten that learning is a much richer and murkier domain. Beyond RL are forms of learning that work without supervision of any kind, that make use of observation, and that require only a single exposure.

### Bayesian decision theory

Decision theory is a branch of mathematical psychology that deals with choosing based on noisy samples. It was inspired by the problem of code-breaking and fueled for many years by the ability to explain human reaction time and performance data across a range of domains^42^. It thus has both a normative basis and empirical support. The diffusion-to-bound model has been linked to single neuron recordings, implemented using the circuit-level dynamic models and derived as a subcase of Bayesian theory. Decision-making is clearly a limited aspect of cognition, but integration is a key process not only in accumulation of evidence, but in working memory, cue combination, and Bayesian inference, etc. Evidence for non-leaky, noiseless integration even in rats^43^ and, the ability to account for other aspect of behavioral data, for example, action timing^14^, contribute to the attractiveness of these models as a basis for describing cognition.

Quantum cognition is an offshoot of decision theory and mathematical psychology that borrows the quantum probability formalism based on Dirac and Von Neumann axioms. Quantum decision models can explain several anomalous effects in sequential decision-making^44^. A quantum diffusion-to-bound model was introduced^45^, which is a promising start toward other cognitive mechanisms. In the background, as either a beacon or a specter, is whether it is ultimately tied to quantum mechanics or quantum computing, as infamously proposed by Penrose^46^.

### Information theory

An information processing metaphor, in which the brain is essentially an input/output system has dominated cognitive science since its origins. Information theory quantitatively formalizes the communication of signals using the fundamental concept of entropy. In neuroscience it has been used successfully to understand how neural signals propagate through sensory systems^47^ and provides a useful basis for conceptualizing the efficiency of neural representations or codes. Moreover, information theory has been used to define what constitutes a “conscious” process^48^,^49^ and recently, it has been shown that simple forms of “cognitive” behavior, such a solving a puzzle or cooperating socially, can emerge from principles of entropy maximization, where future entropy acts as a force in the present^50^. Under the rubric “free energy”, and married with Bayesian and RL theory, such approaches form the core of a proposed a universal brain model^51^.

### Theory of computation

Together with information, the notion of computation has long been a dominant abstraction for conceiving the brain’s most complex functions. Perhaps most fundamentally, Kolmogorov complexity provides a definition of randomness based on the shortest computer program that can be run on a Turing machine^52^. Algorithmic complexity may be useful in application to cognition by offering a grounded definition of what is hard or complex. Despite the dominance of the computational metaphor, there has been surprisingly little contact between cognitive neuroscience and most aspects of computer science. Interestingly, it has been suggested that practical advances in the field of computer engineering (e.g. operating systems) are not themselves well-understood by computer scientists^53^, which may partly explain why current approaches to thinking about cognition remain relatively uninfluenced by advances in computer engineering.

## 6. Machine cognition

The theoretical approaches just outlined are rooted in mathematics and deep concepts—probabilistic inference, learning, entropy, complexity—that are being mined for their potential as an ultimate level of explanation for brain function. These theories have also driven, in parallel, the quest to artificially engineer intelligent machines. In particular, the field of neural networks or machine learning has been closely related to the field of computational neuroscience. What might be the relevance of machine learning, beyond a basis in some of the same normative theories, to defining cognition?

Artificial intelligence (AI) is a practical perspective by which to define what brain functions are truly “hard”, and therefore “cognitive”, under our current theoretical assumptions. It has long been understood that for “symbolic” AI, problems like chess are easy while problems like vision are impossible, not to mention the fact that robots continue to play miserable soccer. Neural networks based on “deep learning” appear to have cracked what had been considered difficult problems such as image recognition. As these approaches are pushed, their limits may come into clearer focus before changing again as new algorithms emerge. Already, hybrid approaches combing, for example, neural networks with Turning machines^54^, are proliferating.

Neural networks share some core features with brains, including distributed processing over simple units and modifiable connections. But they omit the vast majority of what we know about brains, from spikes to cortical layers. And what makes deep learning successful has not been the incorporation of more exacting neural principles but rather clever algorithms such as the back-propagation learning rule that has little to do with brain architecture^55^. It remains uncertain to what degree machine learning approaches will benefit from incorporating more of the brain’s architecture. Consider how divergent are the architectures of the cephalopod and the mammalian brain. Pessimistically (or not), brains and machines might evolve two different solutions to intelligence built on two different sets of tricks, with neither illuminating much the other.

## 7. Cognition informed by behavior

If we get behavior wrong, we will not get cognition right. As we have argued recently^56^ behavior is the foundation of neuroscience, not just a handy tool to investigate the function of the nervous system. When seen as a phenomenon of its own right-not as a mere neural side-effect— the study of behavior becomes invaluable to reveal the nature of cognition.

Bottom up approaches aim to discover, from sophisticated analyses, structure in the copious and high-resolution data currently available. In computational ethology^57^, state-of-the-art approach is the discovery of stereotyped behaviors directly from the images themselves^58^,^59^. Such remarkable data analysis progresses will ultimately need to be incorporated in the context of falsifiable theories. Behavior itself is also a worthy subject of “top-down” theoretical consideration that may inform and perhaps even upend our notion of cognition. Among other potential avenues, one that seems particularly necessary today is to reconsider the idea of negative feedback control, the basis for the “cybernetic revolution” of the 1940’s. Maxwell’s 1867 seminal paper on “governors” described autonomous systems that can maintain certain variables to a reference value despite external disturbances (such as the tray of a waiter as he walks through the restaurant). In fact, negative feedback control is closely related to the concept of homeostasis.

While cybernetic concepts thoroughly transformed engineering practice, they had relatively little impact on the study of behavior. Their potential relevance was however eloquently championed by Powers, who proclaimed that behavior is the control of perception^60^. Then, to understand behavior is to discover the variables that the organism controls, not how it responds to certain stimuli. The implication of this view is that linear cause-and-effect explanations will fail because they assume that real behavior is open loop, when it is not^61^. Behavioral approaches that focus on manipulating the input (stimuli, rewards) and measuring the output (responses, actions) tend to confound controlled with controlling variables. The difference is crucial: the former are controlled by the behaving system whereas the latter are controlled by the human studying the behaving system^62^. One might then be studying the properties of the environmental feedback mistaking them for the behavioral function itself. Note, for instance, the distinction between the behavior of a fan and a thermostat: the first controls its output, the second its input. The thermostat certainly “emits” heat, but that is not really what it does. What it does (or attempts to do) is to specify the temperature to be sensed.

Therefore, rather than concentrating on what you see an animal doing, what is relevant is what the animal is trying to perceive. This double inversion (from my point of view to the animal’s and from action to perception) has three critical behavioral implications for the study of cognition: (i) motor output is a side effect of perceptual control and so mere quantitative data collection data will not suffice, (ii) averaging across individuals may smear out control variables, and (iii) restrained experimental setups may not let animals control the relevant inputs; freedom in the requisite dimensions is required. In short, control (circular causality) and subjectivity (animal centrism) are essential ingredients in behavioral and cognitive neuroscience.

## 8. Cognition beyond the brain itself

Finally, we note two important and relatively new approaches that assert that the data relevant to cognition do not lie solely within the realm of neuroscience facts as traditionally construed. First, “neurophenomenology”, introduced by Francisco Varela, aims to include introspection as a source of scientific evidence^63^. Neurophenomenology is based on first-person (subjective) moment-to-moment accounts coupled with classical quantification procedures such as neuroimaging. Together, these are proposed to provide mutual constraints to tighten the link between neurodynamics and conscious experience and, in particular, to offer new insights in the study of variability and spontaneity in cognition^64^. Second, “embodied cognition”, also championed by Varela, centers on the recognition that cognition is embedded in a context beyond the brain itself, including the body itself as well broader environmental, biological and cultural systems^65^. Both neurophenomenology and embodied cognition illustrate the nascent fruits of substantive dialogue between philosophy and neuroscience that we hope will continue to instill new perspectives in the years to come.

## 9. Conclusions

The neuroscience of cognition is in its infancy. Greatly empowered by new tools, it is still largely hamstrung by the perspectives of the past. In formulating the problems of the future, the field should benefit by paying careful mind not only to neural data but also to the nature of the phenomena to be explained. If the questions we ask are restricted to the same terms they have traditionally been formulated in, potential understanding will be greatly restricted. The various disciplines and frameworks we have summarized here offer rich and widely differing perspectives on how to conceive of cognition. Their mutual incoherence attests to a conceptual vitality that is not merely appropriate but critical to a young field tackling an immense problem. Going forward, it is important for the field to continue to evolve, drawing inspiration from any and all potentially relevant quarters and resisting the tendency to become stuck in a singular, archaic, view of the mind. When we define cognition, we are helping, after all, to define what it is to be human.

**Figure.**
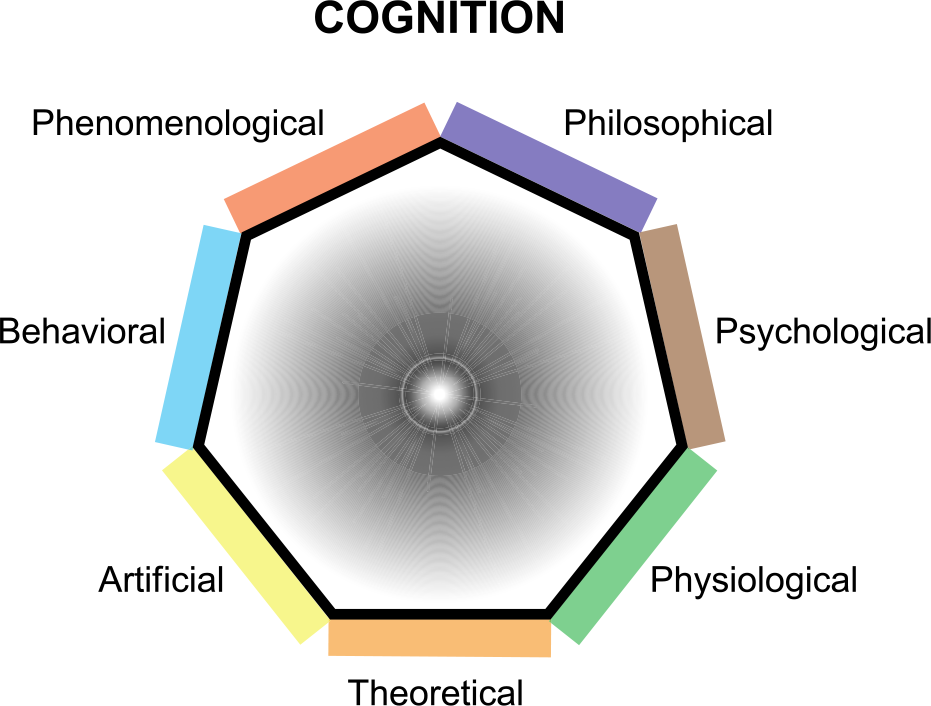
A pluralistic landscape of abstractions to approach the phenomenon of cognition

